# Engineering stringent genetic biocontainment of yeast with a protein stability switch

**DOI:** 10.1101/2022.11.24.517818

**Authors:** Stefan A. Hoffmann, Yizhi Cai

## Abstract

Synthetic biology holds immense promise to tackle key problems we are facing, for instance in resource use, environmental health, and human health care. However, comprehensive safety measures are needed to deploy genetically engineered microorganisms in open-environment applications. Here, we describe a genetic biocontainment system based on conditional stability of essential proteins. We used a yeast-adapted destabilizing domain degron, which can be stabilized by estradiol addition (ERdd). Leveraging the yeast GFP collection and lab automation platforms, we ERdd-tagged 775 essential genes and screened for strains with estradiol dependent growth. Three genes, *SPC110, DIS3* and *RRP46*, were found to be particularly suitable targets. Respective strains showed no growth defect in the presence of estradiol and strong growth inhibition in its absence. *SPC110-ERdd* offered the most stringent containment, with an escape frequency of 7.0×10^-8^, and full growth restoration at 100 nM estradiol. By systematically analyzing the containment escapees, we identified the non-essential C-terminal region of *SPC110* as target for escape mutations. Its removal decreased the escape frequency with a single ERdd tag further to 4.3×10^-9^. Combining *SPC110-ERdd* with a second ERdd tag on either *DIS3* or *RRP46* resulted in escape frequencies below the detection limit of the used assay (<2×10^-10^). Being based on conditional protein stability, this approach is mechanistically orthogonal to previously reported intrinsic biocontainment systems. It thus can be readily combined with other systems, for instance ones based on transcriptional or translational control of essential gene expression, to achieve multiplexed, extremely stringent control over the survival of engineered organisms.

**Significance:** Synthetic biology holds enormous potential to tackle key issues humanity is facing and can for instance revolutionize agriculture, bioremediation or health care. In each case, the unchecked spread of engineered organisms in natural environments must be prevented. This is particularly problematic with use cases of engineered microbes in open environments. Intrinsic, genetically encoded biocontainment systems, which control cell survival based on environmental cues, can solve this issue. We have developed such a genetic biocontainment system acting on the stability of essential proteins, leveraging a switchable degron. Through a large-scale screening for suitable essential target genes, we were able to create yeast strains that are strictly dependent on estradiol. Supplied with this small molecule, the engineered cells maintain high fitness and grow as robustly as the unmodified strains.

## Introduction

Fueled by decreasing cost of DNA sequencing and synthesis, as well as an expanding toolkit for genetic manipulation of organisms from all kingdoms of life, synthetic biology is a rapidly advancing field. It holds promise to tackle key problems humanity is facing, *e.g*. by making agricultural and industrial production more efficient and sustainable, by providing bioremediation of environmental pollution or by offering novel solutions for previously intractable healthcare needs. Comprehensive safety engineering, a hallmark of every mature engineering technology, is required to safely deliver on this promise and pre-empt risks that could emanate from synthetic biology technologies and products.

One layer of biosafety engineering is the development and deployment of genetic biocontainment technologies. It is intended to curb an organism’s ability to survive or transfer its genetic material outside of defined, permissive growth conditions. This intrinsic containment of genetically modified microorganisms is particularly attractive for large-scale or open-environment applications, or when working with pathogens. Historically, genetic biocontainment work has been largely focused either on auxotrophies created by gene knockouts (1, 2), or on control using different ‘suicide genes’, effectors like toxins or Cas endonucleases, with varying degrees of regulatory complexity (3–10). Auxotrophic containment is generally genetically stable but has the caveat that the essential metabolite might be available in certain natural environments, *e.g*. by cross-feeding. In contrast, ‘suicide’ systems typically have a straightforward evolutionary trajectory for escape via the mutational inactivation of the effector gene. Due to these limitations of auxotrophic and suicide containment, control of essential genes for containment presents an attractive alternative approach. Dedicated containment systems targeting essential genes along the flow of genetic information, from genomic presence (11) to transcription (11–13) and translation (13), have previously been developed. Combining different orthogonal control measures allows creating extremely stringent containment (11, 13).

An approach so far not systematically explored for biocontainment is control over the *in vivo* half-life of essential proteins, adding post-translational control over essential genes. There are several switchable degron systems that allow targeted protein degradation in eukaryotes and have been used to link yeast cell viability to cues of light (14), temperature (15) or small molecules (16–18). Control by small molecules absent in most natural environments seems to be an attractive approach to create versatile biocontainment systems. The auxin inducible degron system (16, 17) and the SMASh tag (18) are small molecule-controlled switchable degrons that have been used in yeast on essential proteins, creating conditional cell fitness. Both are functional OFF switches, degrading the target protein in the presence of the molecule. However, this operation logic when applied to essential genes is not suitable for most biocontainment applications, in which proliferation in natural environments is to be prohibited.

Another class of switchable degrons functions as ON switch: the destabilizing domain (DD) degrons (19, 20). Here, we describe the systematic development of a highly stringent containment system based on an estrogen receptor-derived DD degron optimized for yeast (ERdd) (Figure 1A). By screening more than 70% of *S. cerevisiae* essential genes, we have identified particularly suitable targets for biocontainment with this approach. Systematic characterization of containment escape modes allowed engineering an improved system. We further demonstrated the feasibility to multiplex these protein switches and have constructed safeguarded strains with escape frequencies of <2×10^-10^ that maintain wild-type-like fitness.

**Figure 1:**
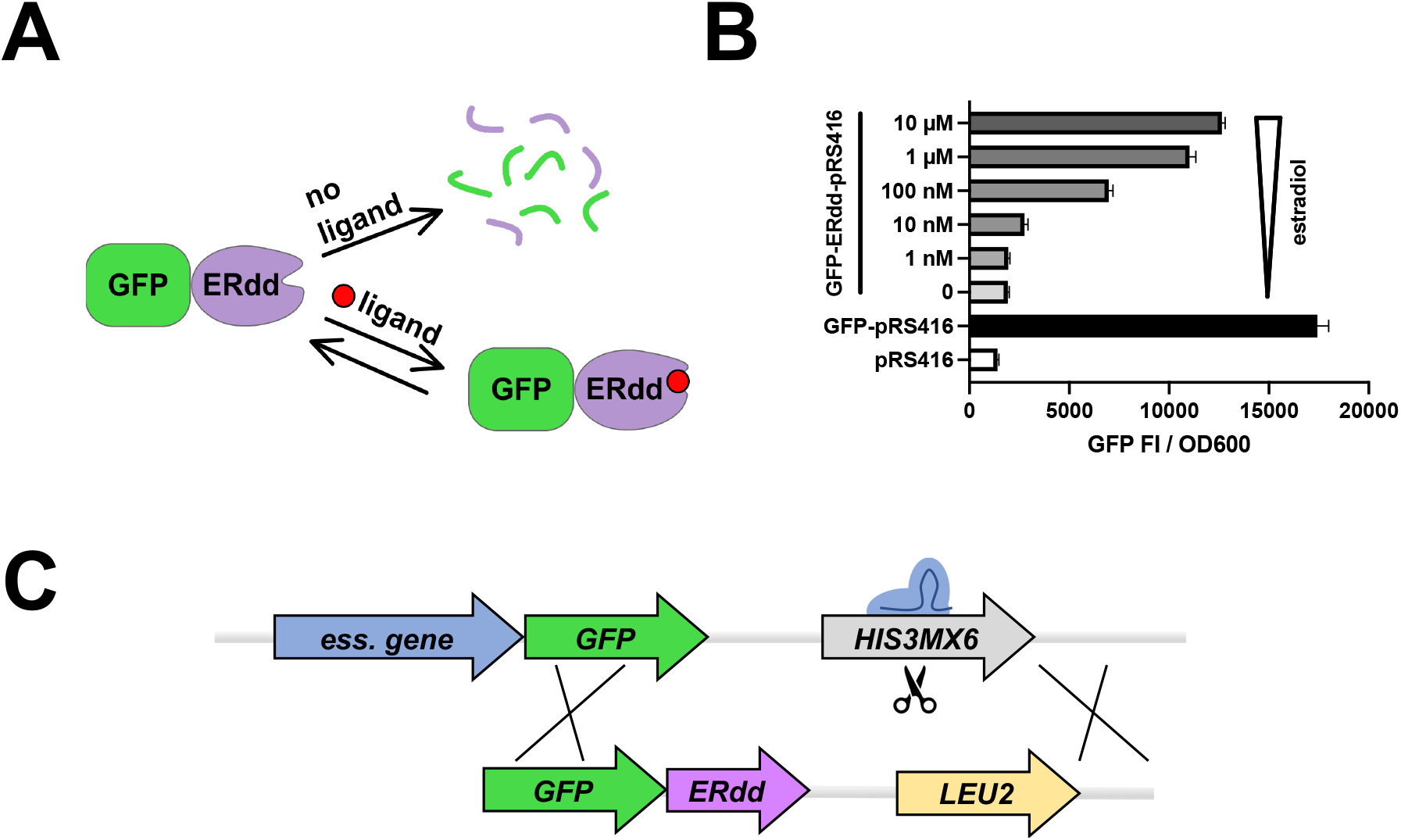
ERdd tag for targeted protein degradation control in yeast. **(A)** In the absence of its ligand, ERdd is in an unordered conformation, triggering its proteasomal degradation along with the protein it is fused to (here shown GFP). Availability of the ligand stabilizes ERdd and greatly increases half-life of the fusion protein. **(B)** Estradiol dependence of GFP fluorescence in yeast with a centromeric plasmid expressing a GFP-ERdd fusion protein. (**C**) Scheme of Cas9-assisted conversion of essential GFP library to essential GFP-ERdd library with switch of marker gene from *HIS3MX6* to *LEU2*.

## Results

### ERdd system in yeast

The *in vivo* behavior of the yeast-adapted ERdd tag was tested by expressing a GFP-ERdd fusion protein from a centromeric vector over a range of estradiol concentrations and assessing GFP fluorescence (Figure 1B). In the absence of estradiol, specific GFP fluorescence was reduced by 95% compared to the fluorescence with 10 μM estradiol. At this highest tested estradiol concentration, specific GFP fluorescence of the GFP-ERdd samples reached 70% of that of untagged GFP. This indicated an exploitable dynamic window to leverage the ERdd tag to couple organism survival to estradiol supplementation by fusion to suitable essential proteins.

### Essential GFP-ERdd library creation and high-throughput screening for estradiol response

We performed a large-scale screen of essential genes for the desired growth response when fused to the ERdd tag – uncompromised fitness in the presence of 1 μM estradiol (permissive condition) and a severe fitness impediment in its absence (restrictive condition). To this end, the yeast GFP collection was leveraged to facilitate ERdd tag integration. The yeast GFP collection contains 4,159 strains, in which ORFs are individually tagged with a C-terminal GFP and a downstream *HIS3MX6* selectable marker gene. This constant region was used as a landing pad for C-terminal addition of the ERdd tag, allowing to use a single donor construct for editing of all strains of interest (Figure 1C). The edit was mediated by a CRISPR/Cas9 system with an integration cassette also swapping the *HIS3MX6* marker for a *LEU2* marker gene, facilitating efficient editing and selection of correct edits. Out of 1103 *S. cerevisiae* ORFs deemed essential^1^, 822 are physically represented as strains in the yeast GFP collection. Out of these, 775 strains (94%) were converted into GFP-ERdd fusion strains (Supplementary Data S1). The others were either missing (24) in the library distribution we had, did not grow (4), or no Leu+/His-colonies could be obtained (19) upon transformation of the CRISPR/Cas components.

From the constructed essential GFP-ERdd library, estradiol dependent strains were identified by a bipartite screening (Supplementary Data S2). The primary screening for estradiol response was performed by pinning the generated essential GFP-ERdd library on YPD without (restrictive) and with 1 μM estradiol (permissive) and analyzing the colony size ratios (Figure 2A). For most of the essential genes, ERdd-dependent degradation without estradiol appeared to not be substantial enough to cause a growth-inhibition phenotype, but about 5% of the screened strains showed a clear dependence on estradiol for proper growth (Figure 2B). The 46 strains with the highest ratio of colony size between permissive and restrictive conditions were chosen for a subsequent screening in liquid culture in plate reader assays. For almost all of these strains differential growth in response to estradiol was observed in the liquid culture screening as well (Figure 2C,D). Strains were ranked by the mean area under the difference curve between growth under permissive and restrictive conditions.

**Figure 2:**
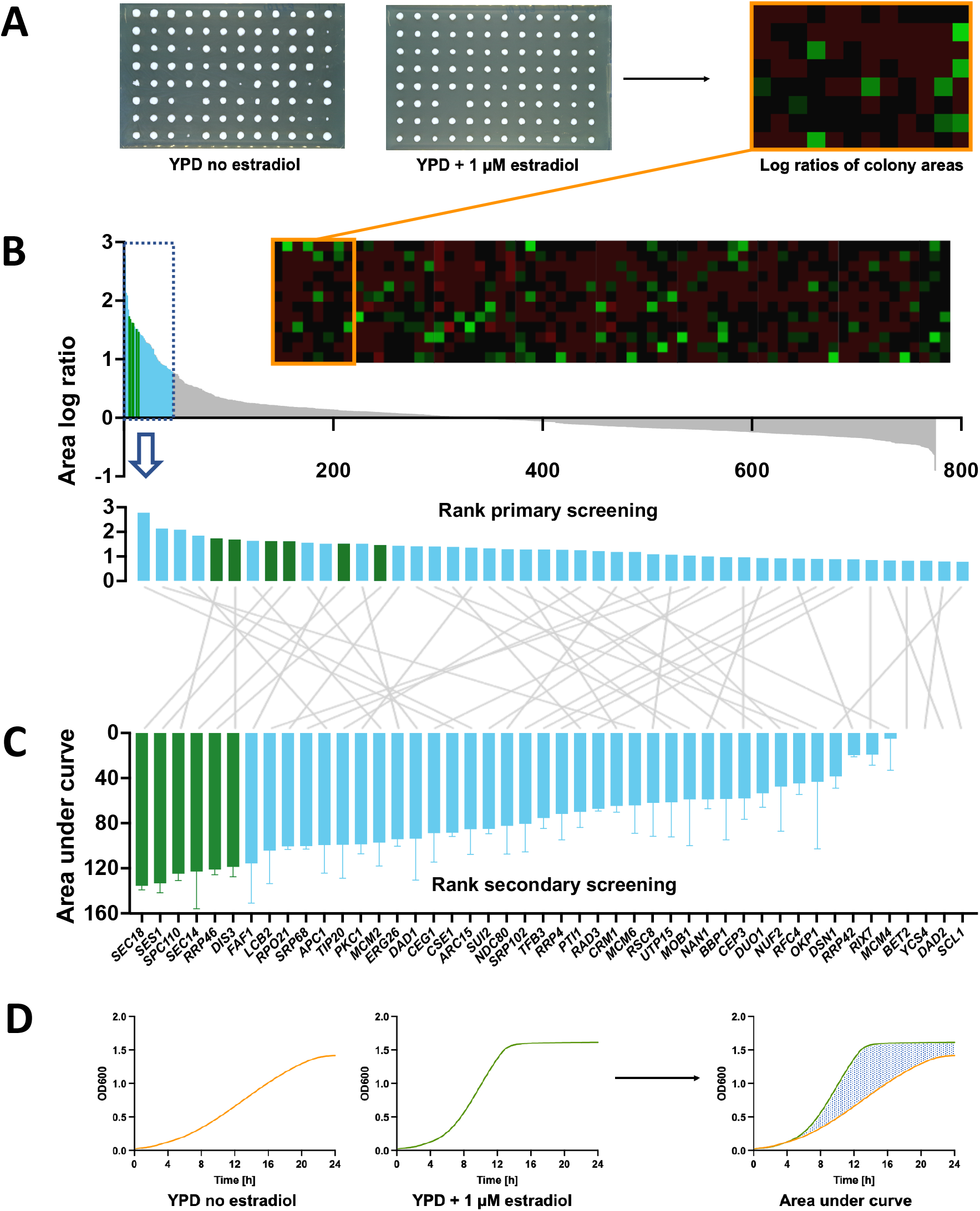
Bipartite screening for estradiol dependence of essential GFP-ERdd strains. (**A**) For the primary screening, essential GFP-ERdd strains were pinned on YPD agar without and with 1 μM estradiol. The area log ratios between permissive and restrictive condition were determined and used as metric for ranking. In the heatmap bright green indicates a strongly positive log ratio, *i.e*. a larger colony area with estradiol than without. (**B**) Measured area log ratios for the 775 strains in the primary screening were ranked from largest to smallest and the 46 highest-scoring strains were subjected to (**C**) the secondary screening in liquid culture. The 6 highest-scoring strains in the secondary screening (marked in green) were created as direct fusions in a BY4742 background. (**D**) In the secondary screening, growth curves with and without estradiol were recorded for the 46 select strains, and the area of the difference curve was used as metric for ranking.

Most of the screened essential gene-*GFP-ERdd* fusions did not show a clear estradiol-dependent phenotype. To investigate whether this was due to a lack of ERdd-mediated essential protein degradation, the GFP-ERdd strains of the highly expressed essential genes *PGK1, YEF3, TPI1* and *GPM1* were assayed for growth and GFP fluorescence when cultured with and without estradiol (Figure S1). Relative fluorescence intensities suggested that substantial proportions of the respective fusion proteins were being degraded upon estradiol withdrawal (47-82% at 24h). However, evidently the remaining essential protein amounts were sufficient to support growth, as the strains showed either no or only minor growth inhibition in the absence of estradiol.

### Estradiol response of single and double ERdd tagged strains

The genes corresponding to the six highest ranked strains in the secondary screening, *SEC18, SES1, SPC110, SEC14, RRP46* and *DIS3*, were directly fused to ERdd in a BY4742 background without using a marker gene. The resulting six strains all showed a strong dependency on estradiol for growth (Figure 3A, Figure S1). A genetic safeguard should not compromise the cell fitness under permissive conditions, as to not create an evolutionary incentive to inactivate the safeguard. Three of the strains, *SPC110-ERdd, RRP46-ERdd* and *DIS3-ERdd*, exhibited a growth indistinguishable from the parental strain in permissive medium (YPD with 1 μM estradiol) (Figure 3B) and were used for further work. Of these, *SPC110-ERdd* displayed particularly favorable behavior, with a severe growth inhibition without estradiol and full restoration of growth already at 100 nM estradiol (Figure 5A). Fusion of ERdd to a truncated *SPC110*, removing its non-essential C-terminal region, resulted in a strain with similarly favorable estradiol dependence; growth of *SPC110D845-ERdd* likewise was fully restored at 100 nM estradiol (Figure 5C).

**Figure 3:**
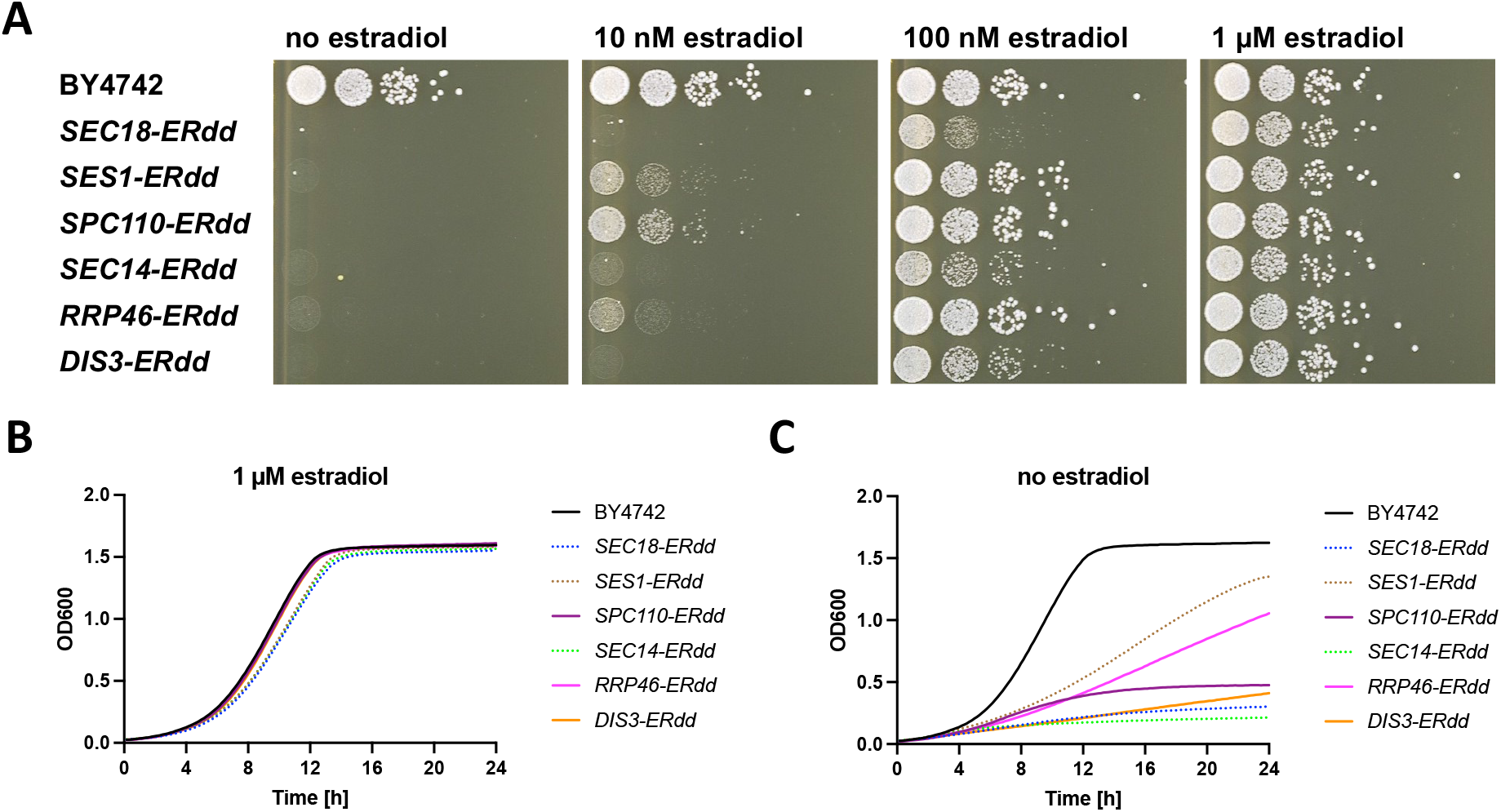
Assaying growth of ERdd fusion strains for estradiol dependence. (**A**) Spot assay of strains with direct ERdd fusion to a single essential gene at different estradiol concentrations on YPD agar, showing strong dependence on estradiol for all tested ERdd strains. (**B**) Growth assay in YPD with 1 μM estradiol, showing strains with growth indistinguishable from that of the parental strain BY4742 in solid lines and ones with a slight apparent growth defect with dashed lines. (**C**) All tested ERdd strains show a substantial to very strong growth defect without estradiol.

Subsequently, strains with two ERdd-tagged genes were created for combinations of *SPC110, RRP46* and *DIS3*. All dual ERdd strains exhibit strict dependence on estradiol. Growth of *SPC110-ERdd/RRP46-ERdd* was fully restored with 100 nM estradiol, whereas the other two strains required a higher estradiol concentration, reaching wild-type-like growth at 1 μM estradiol (Figure 4, Figure S2).

**Figure 4:**
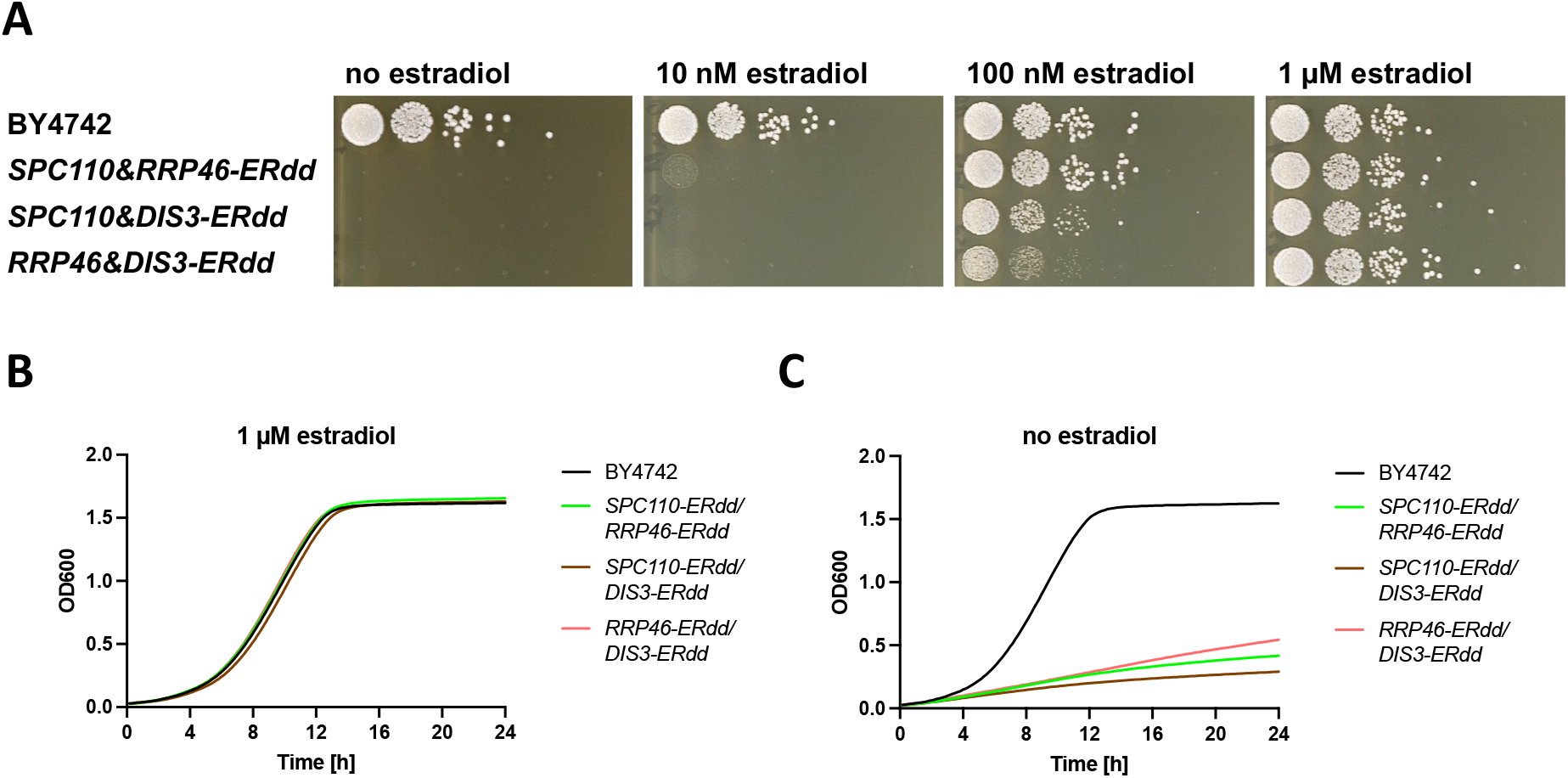
Assaying estradiol dependence of strains with two essential genes fused to ERdd. (**A**) Spot assay of strains at different estradiol concentrations on YPD agar, showing strict dependence on estradiol for all dual ERdd strains. Growth assay of dual ERdd strains in YPD (**B**) with 1 μM estradiol and (**C**) no estradiol.

### Escape rates and analysis of escapees

The estradiol regulated strains not exhibiting any growth impairment under permissive conditions, *SPC110-ERdd, RRP46-ERdd, DIS3-ERdd* and derived double ERdd-tagged strains, were assessed for the stringency of containment they offer. This was done by assaying the frequency of escape colonies occurring on restrictive medium. To this end, eight independent cultures of the assayed strain were plated on YPD without estradiol (restrictive environment) and at a suitable dilution on YPD with estradiol (permissive environment) to determine the number of colony-forming units.

Afforded containment varied drastically between the three genes when individually tagged with ERdd. *SPC110-ERdd* had the lowest escape frequency of 7.0×10^-8^, followed by *DIS3-ERdd* with 6.1×10^-7^. *RRP46-ERdd* kept growing very slowly on restrictive plates, leading to the continued emergence of escaper colonies with prolonged incubation, making the quantification of an escape frequency infeasible. Combining *SPC110-ERdd* with an ERdd tag on either *RRP46* or *DIS3* resulted in stringently contained strains with escape frequencies below the detection limit of this assay, not yielding a single escapee colony. Subsequently, assays for low escape frequencies were carried out for these strains, plating about 5× 10^9^ colony forming units (CFUs) on restrictive plates, which were replica plated for single colony detection. Again, no escapee colonies were observed. Accordingly, escape frequencies of *SPC110-ERdd/RRP46-ERdd* and *SPC110-ERdd/DIS3-ERdd* were below the assay’s detection limit of 2×10^-10^. In contrast, ERdd tagging both *RRP46* and *DIS3* yielded an escape frequency of 4.3×10^-7^, not markedly improving containment over singly tagged *DIS3-ERdd* (6.1×10^-7^). Interestingly, both *RRP46* and *DIS3* code for a subunit of the exosome complex essential for RNA metabolism.

Eight escapee strains from each of the three singly ERdd-tagged were isolated, and the ERdd-tagged gene was Sanger sequenced. For the sequenced *SPC110-ERdd* escapees, seven unique escape mutations were found, mapping to the C-terminal part of the Spc110 protein or the linker to the ERdd tag (Figure 5). All of them result in a premature stop, leading the absence of the ERdd tag. Seven of the eight mutants were missing a C-terminal portion of the protein.

**Figure 5:**
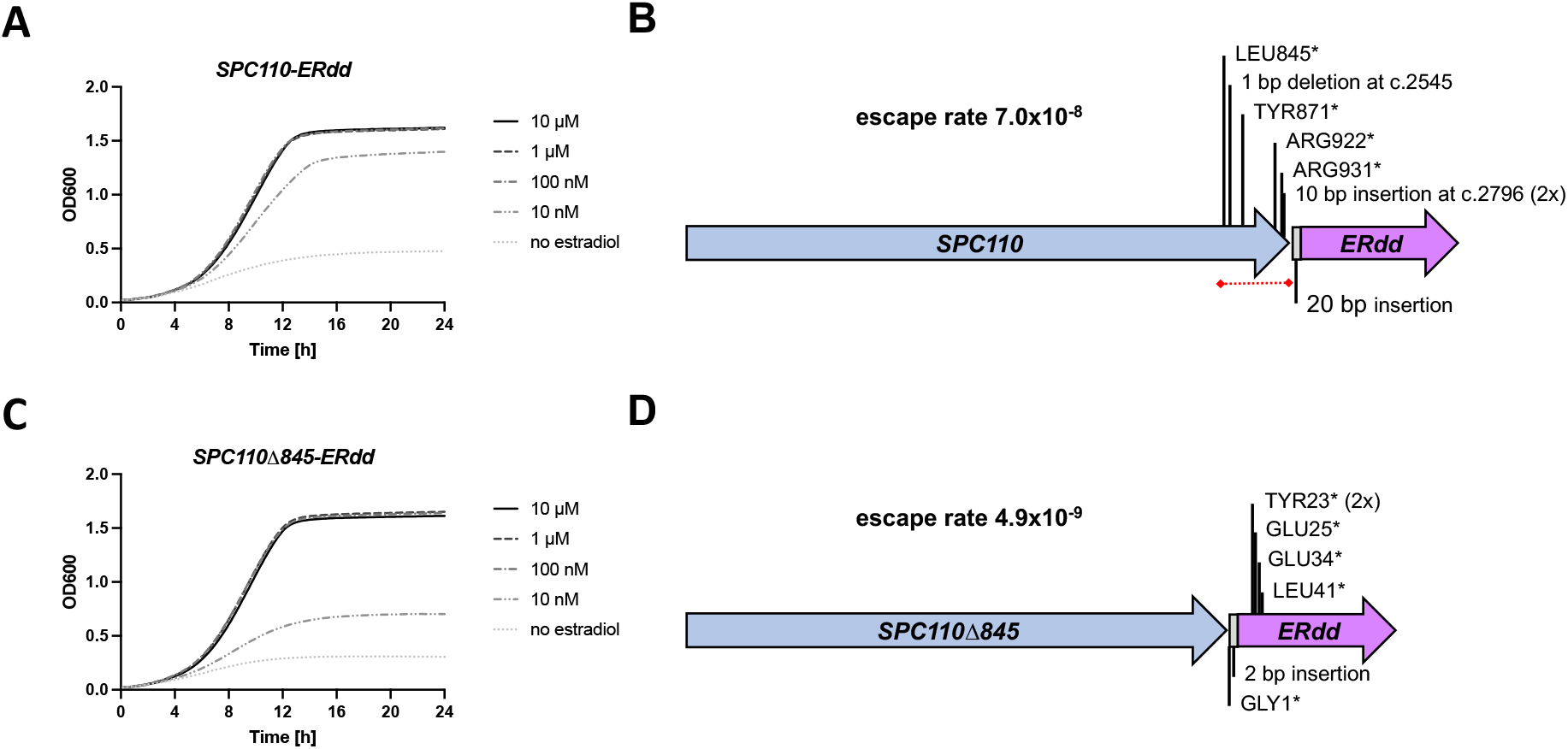
Escapee analysis of *SPC110-ERdd* and engineering for improved containment stringency. (**A**) Growth of *SPC110-ERdd* at different estradiol concentrations in YPD. (**B**) *SPC110-ERdd* containment escape mutations. Eight *SPC110-ERdd* escapees were sequenced, and seven unique escape mutations were found. All resulted in a premature translation stop by mutation to a stop codon or a frameshift, mapping to the C-terminal region of *SPC110* or the linker, and conversely removal of the ERdd tag. No mutations were found in *ERdd* itself. The dispensable C-terminal region of *SPC110* in which escape mutations clustered is marked with a red dashed line. (**C**) Growth of *SPC110Δ845-ERdd*, in which the C-terminal region has been removed, at different estradiol concentrations in YPD. As with *SPC110-ERdd*, growth is fully restored by 100 nM estradiol. (**D**) *SPC110Δ845-ERdd* containment stringency is markedly improved over *SPC110-ERdd*. Seven *SPC110Δ845-ERdd* escapees were sequenced. Each contained a unique escape mutation, two of which resulted in a stop codon at the same position (TYR23* in ERdd). All of the sampled *SPC110Δ845-ERdd* escape mutations mapped to the linker or the ERdd tag.

The C-terminal region of Spc110p is known to be involved in binding to calmodulin, but not essential for viability (21). Interestingly, no mutations within the ERdd tag itself were found, indicating its resistance against escape by truncation. Based on this finding, we generated an ERdd fusion omitting the non-essential C-terminal domain, *SPC110Δ845-ERdd*. This indeed increased containment stringency without compromising growth under permissive conditions. The strain’s escape frequency was below the detection limit of the standard escape assay, not yielding escape colonies. In a low escape frequency assay an escape rate of 4.3×10^-9^ was determined. All of the sampled *SPC110Δ845-ERdd* escape mutations mapped to the linker or ERdd tag, demonstrating that escape routes through truncating mutations in *SPC110* itself have been closed off by removal of its C-terminal region.

The sampled *RRP46-ERdd, DIS3-ERdd*, and doubly tagged *RRP46-ERdd/DIS3-ERdd* escapees had no escape mutations in the respective fusion genes. To elucidate their escape mechanism, the whole genome of three clones of each were sequenced by Illumina sequencing (Supplementary Data S3). For each of the six escapees with an ERdd-tagged *DIS3* gene, a mutation likely to reduce proteasome activity could be found. Three unique mutations (two non-sense, one frameshift) of *RPN4*, a transcription factor stimulating expression of proteasome genes were sampled. Further, a non-sense mutation of a chaperone for proteasome maturation (*UMP1*), and non-synonymous mutations of two proteins involved in proteasome regulation, *RPN3* and *RPT5*, were found. Likewise, for one of the *RRP46-ERdd* escapees a non-synonymous mutation of a proteasome subunit was the likely cause for containment escape. For the other two *RRP46-ERdd* escapees no obvious suppressor mutations could be identified.

### Competitive fitness of contained strains

The perfect containment system imparts no fitness defect under permissive conditions as to not provide an evolutionary incentive to inactivate the containment system under prolonged culture. The strains contained by single or dual ERdd fusion to *SPC110*, *DIS3*, or *RRP46* and respective combinations showed growth that was indistinguishable from that of the parental BY4742 with 1 μM estradiol in growth assays. However, a much more stringent measure for fitness is the competition against the parental strain under repeated, lab-typical batch cultures, cycling through different physiological states. Competition cultures were carried out over ten such cultures cycles for a total of 100 generations in YPD with 1 μM estradiol (Figure 6A), assessing the frequency of the parental and safeguarded strain at 0, 50 and 100 generations from 46 to 48 samples each. None of the above-mentioned single and dual ERdd strains appeared to be outcompeted by the WT parental strain. This apparent lack of evolutionary incentive for containment inactivation should result in a high stability of the containment system in prolonged culture under permissive conditions. This was directly assessed by cultivation of eight independent *SPC110-ERdd* cultures over 100 generations under permissive conditions followed by assaying the escape frequencies. The measured average escape frequency after 100 generations was 2.2×10^-8^, not having increased following prolonged culture.

**Figure 6:**
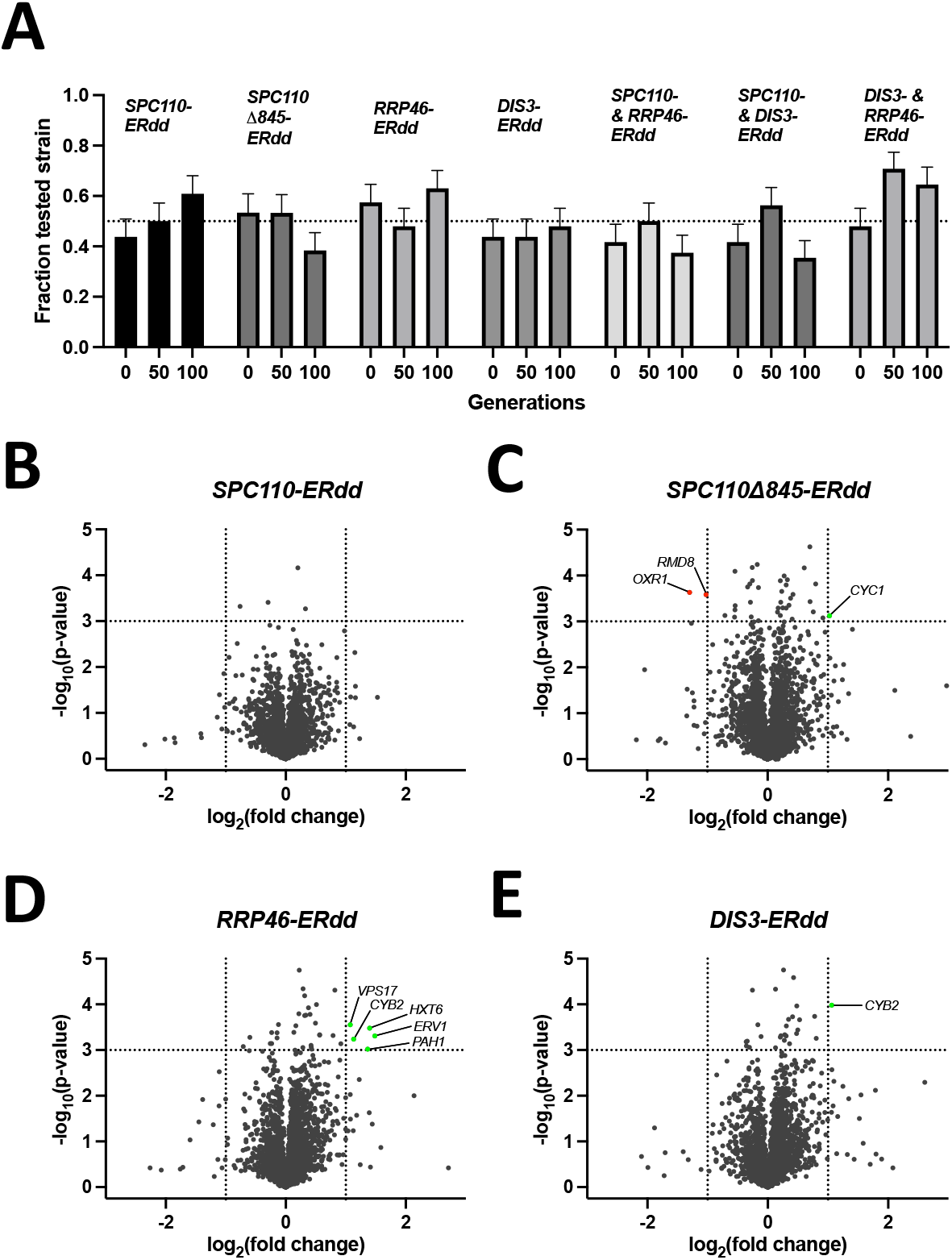
Competitive fitness and physiological characterization of contained strains **(A)** Competition of singly and doubly ERdd-contained strains against BY4742 under permissive condition, in YPD with 1 μM estradiol. Respective contained strains were mixed 1:1 with BY4742 and grown for a total of 100 generations, with the relative frequencies of respective strains assessed after 0, 50 and 100 generations by colony PCR. **(B)** to **(E)** show volcano plots of relative normalized protein abundances measured by LC-MS compared to the parental strain, using raw p-values of pairwise comparisons.

### Proteome analysis of contained strains

To investigate effects of the addition of ERdd on cell physiology under permissive conditions, cells grown with 1 μM estradiol were harvested in mid-log phase and analyzed by LC-MS. Normalized protein abundances of the safeguarded strains were compared to those of the parental strain. Between 2939 and 2943 protein abundances could be compared for each of the strains. For *SPC110-ERdd* no significant perturbances of the proteome were detected, whereas *SPC110Δ845-ERdd* (1 up-, 2 down-regulated), *RRP46-ERdd* (5 upregulated) and *DIS3-ERdd* (1 upregulated) showed apparent detectable changes of the proteome. Interestingly, in both *RRP46-ERdd* and *DIS3-ERdd* cytochrome b2 appeared to be upregulated. Both *RRP46* and *DIS3* are components of the RNA exosome complex, but the relationship to cytochrome b2, a mitochondrial protein involved in lactate utilization, is unclear.

## Discussion

The availability of stringent, stable and use case-suited intrinsic biocontainment systems is vital for a next-generation bioeconomy, one that leverages the possibilities of synthetic biology in open-environment applications. Due to their high replicative potential and involved large number of individual organisms, safely and effectively containing engineered microorganisms through genetic systems is particularly challenging. An NIH guideline for work with synthetic or recombinant nucleic acids in laboratory settings calls for systems with an escape rate of less than 10^-8^. Since a single milliliter of a densely grown culture can easily contain 10^8^ cells, this threshold is clearly insufficient for applications on industrial scales. Reaching adequate containment stringency including a generous safety margin likely will require the combination of multiple independent systems.

Here, we have developed a new class of genetic biocontainment systems based on conditional stability of essential proteins. Using a destabilizing domain (DD) conditional degron and applying a systematic, large-scale search for suitable essential genes from 775 genes, we have identified *SPC110, DIS3 and RRP46* as suitable target genes for this system. Fusing *ERdd* to any of these genes rendered cells strictly dependent on estradiol without imposing a fitness defect when it is supplied. *SPC110-ERdd* was particularly suited, with a containment stringency under restrictive conditions nearly satisfying the mentioned NIH guideline. Escapee analysis revealed that premature translation stops in the C-terminal region of Spc110p were the most frequent escape mechanism. Fusion of an accordingly truncated *SPC110* to the ERdd tag resulted in escape frequencies of less than 10^-8^. Also, combining *SPC110-ERdd* with an ERdd tag on either of the other two genes resulted in containment greatly exceeding these guideline requirements.

*SPC110-ERdd, SPC110Δ845-ERdd* and *SPC110-ERdd/RRP46-ERdd* required as little as 100 nM estradiol to achieve growth indistinguishable from the wild type. This equates to cost of less than $1 to treat 1000 L of culture, making it a highly cost-effective approach. The systems can be for instance combined with estradiol-dependent transcription of essential genes, which would allow controlling two independently working genetic containment systems with a single, near-gratuitous ligand at nanomolar concentrations. We have previously employed a chimeric, estradiol-inducible transcriptional activator of *GAL* promoters (GAL-EBD-VP16) (22) with select essential genes to create stringent biocontainment (11, 12). Alternatively, a more recent, similar system (ZEV) (23) allowing estradiol-dependent transcription from synthetic promoters could be used. Recently, a comprehensive screen of over 1000 essential genes with the ZEV system was reported, identifying several hundred genes with estradiol-dependent growth when under its control (24). These could be purposefully screened for suitability for biocontainment.

The work presented here shows conclusively that conditional stability of select essential proteins can be leveraged to create stringent and stable biocontainment with specific ligand dependence. However, alternative small molecules might be beneficial use cases in which the utilization of estradiol for biocontainment is undesirable. For instance, large-scale use of estradiol with discharge into the environment can be environmentally problematic due to its endocrine activity (25). Notably, there are more ‘destabilizing domain’ (DD) systems based on different protein scaffolds available that respond to other ligands: DD systems responding to the synthetic ligand Shield-1 (19), trimethoprim (26), and bilirubin (27), respectively, have previously been developed. These alternative DD degrons should allow analogously creating biocontainment systems responding to their respective ligands.

## Materials and methods

### Strains, plasmids, synthetic gene fragments and oligonucleotides

The yeast GFP collection (28) and the *S. cerevisiae* strain BY4742 were used for the creation of the essential GFP-ERdd library (Supplementary Data S1) and generation of direct ERdd fusion strains (Table S1), respectively. DNA constructs made in this study as plasmids were cloned and amplified in *E. coli* DH5α. For creation of gRNA plasmids, pairs of oligonucleotides containing the respective target sequence and bases to create required overhangs (Table S2) were annealed. Subsequently, they were cloned into the gRNA entry vector pWS082 in a Golden Gate reaction (30 cycles of 1 min at 37°C, 1 min at 16°C and heat inactivation for 5 min at 80°C) with 40 nM of annealed insert, 50 ng of pWS082, 5U Esp3I and 20U T4 DNA ligase in 10 μl T4 DNA ligase buffer. The *ERdd GFP LEU2* donor plasmid SHe146 (Supplementary Data S4) was cloned by Golden Gate assembly from parts PCR amplified from pScDD2_ERdd (Addgene plasmid #109047), pWS082 (Addgene plasmid #90516), genomic DNA from the yeast GFP collection, and pRS415 (29). Donors for direct ERdd fusions were ordered as gene fragments from Twist Bioscience (Supplementary Data S5). For confirmation of edits and for strain identification by colony PCR, pairs of primers spanning the end of the respective coding sequence were used (Table S2).

### Creation of essential GFP-ERdd library

The yeast GFP collection (in a *S. cerevisiae* BY4741 background) was spotted in 384 well format on Synthetic Complete (SC) -His agar. The available GFP strains were cross-referenced with a list of *S. cerevisiae* essential genes (essglist). The respective strains with GFP labelled essential genes were cherrypicked onto SC -His agar in 96 well format using a Singer Instruments PIXL colony picking robot. The GFP cassette with the *HIS3MX6* marker gene was used as a landing pad for CRISPR/Cas9-based insertion of the ERdd as a C-terminal fusion to the GFP, while swapping *HIS3MX6* for a *LEU2* marker gene. For transformation, strains were inoculated into YPD medium in a deepwell plate and grown over night at 30°C at 850 rpm. The next day, new cultures were set up in a deepwell plate by inoculating 1 ml YPD with 20 μl of the respective overnight culture each and grown for 4h at 30°C at 850 rpm. The cultures then were pelleted and washed with 1 ml, followed by another wash step with 0.5 ml 100 mM LiOAc. Supernatant was removed and a transformation mixture was added to achieve final concentrations of 33.3% PEG 3350, 100 mM LiOAc, 0.3 mg/ml boiled herring sperm DNA (Promega) with about 200 ng pWS174 digested with Esp3I (Cas9 plasmid), 400 ng of the *HIS3MX6* guide plasmid digested with EcoRV and 1300 ng SHe146 digested with NotI (*GFP-ERdd LEU2* donor). Digested plasmids were purified by isopropanol precipitation and ethanol wash prior to transformation. The deepwell transformation plate was thoroughly vortexed and incubated 1h at 37°C at 850 rpm followed by an incubation at 42°C for 20 min. Cell suspensions were pelleted, the supernatant removed, and the cells resuspended in 950 μl SC - Leu with 1 μM estradiol. These cultures selective for genomic insertion of the donor were incubated at 30°C at 850 rpm. After 18h, nourseothricin was added to a final concentration of 100 μg/ml to select for presence of the reconstituted Cas9 plasmid. After 48h the transformation cultures were pinned onto SC-Leu agar with 1 μM estradiol using a Singer Instruments ROTOR replicating robot in a 7 by 7 pattern for each culture, creating a dilution series for each transformation outgrowth to obtain separated colonies. Up to four individual colonies were picked from each pinned transformation first onto SC-His, then onto SC-Leu and finally onto SC-Leu with 1 μM estradiol agar with the PIXL robot.

### Screening of GFP-ERdd library

The primary screening of the essential GFP-ERdd library for estradiol regulation was done on agar. From the SC-Leu +1 μM estradiol agar plates from the transformation, one (out of up to four) colony from each transformation was spotted with the PIXL robot, first onto YPD and then onto YPD +1 μM estradiol agar in a 96 well layout. Only His auxotroph colonies were picked, and colonies with apparent regulation by estradiol were preferentially selected. The plates were incubated for 2 days at 30°C and then imaged with a Singer Instruments Phenobooth, and colony sizes were assessed. The log ratio of the colony size with estradiol to the size without estradiol was the metric used to select the 46 highest scoring strains for the secondary screening.

The secondary screening was done in liquid culture in a plate reader. The strains selected from the primary screen were grown overnight in YPD with 1 μM estradiol at 30°C and 850 rpm in a deepwell plate. The next day, the cultures were washed once with YPD without estradiol. In a 96 well F-bottom microtiter plate with lid, for each strain a sample of 200 μl YPD without and a sample of YPD with 1 μM estradiol was set up and inoculated 1:100 with the respective washed cell suspension. The assay was run for 24h at 30°C in a BioTek Synergy H1 plate reader, recording the optical density at 600 nm every 10 min while shaking between the measurement cycles. The growth curve without estradiol was subtracted from the growth curve with estradiol and the area under the difference curve was calculated. Strains were ranked by the mean area under the curve averaged from three assay runs.

To investigate degradation of the essential GFP-ERdd fusion protein, growth assays in a plate reader were carried out like the secondary screening, but in SC-Leu and measuring the GFP fluorescence (excitation 479 nm, emission 520 nm, gain 100, with background subtraction) in addition to optical density.

### Markerless ERdd insertion

The six highest ranked candidate genes from the secondary screening were created as direct markerless ERdd fusions in a BY4742 background using a CRISPR/Cas9 approach. A standard LiOAc yeast transformation protocol was used: exponentially growing yeast cells were harvested, washed first with water and then with 100 mM LiOAc. The transformation mix contained cells from 5 ml culture in 400 μl with final concentrations of 33.3% PEG 3350, 100 mM LiOAc, 0.3 mg/ml boiled herring sperm DNA (Promega) and about 500 ng pWS174 digested with Esp3I (Cas9 plasmid), 600 ng of the respective guide construct digested with EcoRV and 1000 ng of the respective donor DNA (Twist Bioscience). After thorough mixing, transformation cultures were incubated for 60 min at 30°C at 200 rpm and then for 20 min at 42°C. Cultures were spun down and resuspended in 500 μl YPD with 1 μM estradiol. After a transformation outgrowth period of 1-2 h at 30°C under shaking, 100 μl were plated on YPD agar with 100 μg/ml nourseothricin and 1 μM estradiol and incubated for 3d at 30°C. Colonies were picked with a PIXL robot onto YPD with 1 μM estradiol. Insertion was confirmed by colony PCR. A correct clone was restreaked both on YPD with 1 μM estradiol, and loss of the Cas9 plasmid was confirmed by spotting colonies onto YPD with 1 μM estradiol and YPD with 1 μM estradiol and 100 μg/ml nourseothricin. Strains with two ERdd fusions were created the same way, starting with strains that already had one of the two genes ERdd tagged.

### Assaying estradiol response

The growth response of individual strains to estradiol was assessed both in liquid culture using a plate reader and in spot assays on solid media. Strains were cultured overnight in YPD with 1 μM estradiol, washed once in YPD, and adjusted to an OD 0.1 in YPD. With these cell suspensions, assays in the plate reader were carried out like the secondary screening, but at 10 μM, 1 μM, 100 nM, 10 nM and no estradiol, with duplicates for each estradiol concentration. For spot assays, log10 dilution series of the OD 0.1 cell suspensions to OD 10^-7^ were prepared and 5 μl of each dilution were spotted on YPD agar at the different estradiol concentrations used (10 μM, 1 μM, 100 nM, 10 nM and no estradiol), incubated at 30°C and imaged with a Phenobooth after two and three days.

### Escape rate analysis

For assessment of escape rates of ERdd strains, 8 individual colonies were picked into YPD with 1 μM estradiol and incubated overnight at 30°C and 850 rpm. The next day the cultures were washed twice with YPD with no estradiol and a log10 dilution series to 10^-5^ was prepared. From each replicate 100 μl of the washed undiluted culture were plated on YPD agar without estradiol, and 100 μl of the 10^-5^ dilution were plated on YPD agar with 1 μM estradiol. Plates were incubated for 2 days at 30°C. The counted number of escapees on restrictive medium (no estradiol) was divided by total colony forming units extrapolated from the colony number on the permissive medium (with estradiol). For each strain the average escape rate was determined from the 8 replicates.

For assessment of low escape frequencies, 50 ml of overnight cultures (in YPD with 1 μM estradiol) were washed twice with water and resuspended in 5 ml YPD. This suspension was plated onto ten Petri dishes with YPD agar without estradiol. After 2 days at 30°C, the dishes were replica plated onto fresh YPD agar without estradiol and incubated for another 2 days at 30°C. To estimate the total plated number of colony-forming units, 100 μl of a 10^-6^ dilution of the washed and concentrated cell suspension were plated on permissive medium.

### Whole genome sequencing

Escapee strains and the parental strain BY4742 were grown overnight in permissive medium (YPD with 1 μM estradiol). DNA was isolated from 1.5 ml of culture with a MasterPure Yeast DNA Purification Kit and taken up in 50 μl TE buffer. To remove RNA contamination of the DNA preparation, 1 μl of 4 mg/ml RNase A (Promega) was added and the sample incubated for 30 min at 37°C. The DNA was then column purified and eluted with 50 μl water. Sequencing library preparation, Illumina sequencing, mapping of reads and variant calling was performed by Eurofins Genomics. Reads were pre-processed with fastp v0.20.0 (30) and mapped with BWA v0.7.17 (31). Subsequent variant calling was done using Sentieon’s HaplotypeCaller. Variants in coding sequences that a) passed quality control filters, b) were not found in the parental strain and c) were present in at least half of respective reads were annotated using VariantAnnotation v1.40 (32).

### Competition and stability assay

For assessing long-term competitive fitness compared with the parental strain, BY4742 and the ERdd strain to be assayed were each picked into YPD with 1 μM estradiol and incubated overnight at 30°C. The overnight cultures were mixed 1:1 and the mixture diluted 1:1000 in YPD with 1 μM estradiol and incubated for 24h at 30°C under shaking. Thereafter, the culture was rediluted 1:1000 for another 24h of growth. This was repeated for a total of 10 days of growth of the mixed culture, representing about 100 generations. Aliquots of the initial mixture and the mixed culture after days 5 and 10 were plated on YPD agar with 1 μM estradiol to assess the genotype frequencies by colony PCR. To investigate long-term stability of the containment of *SPC110-ERdd*, the strain was similarly cultured over 10 days for about 100 generations in 8 independent cultures, from which an escape rate analysis was performed.

### Proteome analysis

Four replicates of containment strains and the parental strain BY4742 were grown overnight in permissive medium (YPD with 1 μM estradiol). Dense overnight cultures were diluted back 1:50 with YPD with 1 μM estradiol and grown for 4h at 30°C. From each culture, 1 ml samples were pelleted and washed twice with PBS. Cells were lysed in 5% SDS and 50 mM TEAB pH 7.5 using a Covaris LE220+ sonicator. Protein lysates were reduced, alkylated, acidified with phosphoric acid and the protein bound to wells of an S-Trap plate. After a tryptic digest, peptides were eluted and desalted using Corning FiltrEX desalt filter plates and OLIGO R3 beads. The liquid chromatography was performed on a Waters nanoEase M/Z Peptide CSH C18 Column using a Thermo RSCL system with a multistage gradient using 0.1% formic acid in water as buffer A and 0.1% formic acid in acetonitrile as buffer B with a runtime of 60 min. Mass spectrometry of the eluted peptides was carried out with a Thermo Exploris 480. Analysis of MS spectra was performed with Proteome Discoverer v2.5.0.400 using NCBI txid559292 v2022-04-30 (*Saccharomyces cerevisiae* S288C) as reference for identification of proteins. Proteins with at least two unique peptides were considered for quantitative comparison. For each analyzed contained strain, normalized abundances of found proteins were compared to the corresponding abundances in the parental strain. Proteins with a raw p-value of <0.001 and a fold change of >2 or <0.5 were considered significantly changed relative to the wild type.

## Acknowledgements

This work is supported by grants from the EPSRC (EP/V05967X/1), the BBSRC (BB/W014483/1) and the Volkswagen Foundation “Life? Initiative” (Ref. 94 771). We thank Shuangying Jiang and Junbiao Dai for providing, and Ellie Payne and Reem Swidah for archiving the yeast GFP collection. We also thank Raymond Wan for variant annotation of whole genome sequencing data, and Elisa Barrow Molina for help with testing degron systems. Further, we thank David Knight and colleagues at the University of Manchester BioMS Core facility (RRID:SCR_020987), in particular Ronan O’Cualain for help with proteomics sample preparation, Stacey Warwood for running the samples, and Julian Selley for bioinformatic analysis. Finally, we also want to thank Thomas Wandless and Rishi Rakhit for helpful discussions on the ERdd system. The Cas9 expression plasmid pWS174 (Addgene plasmid #90961) and the gRNA entry vector pWS082 (Addgene plasmid #90516) were gifts from Tom Ellis. The plasmid pScDD2_ERdd (Addgene plasmid #109047) was a gift from Thomas Wandless.

## Data Availability

Whole genome sequencing data has been deposited at SRA with the BioProject number PRJNA923142.

## Supporting Information

**Figure S1:**
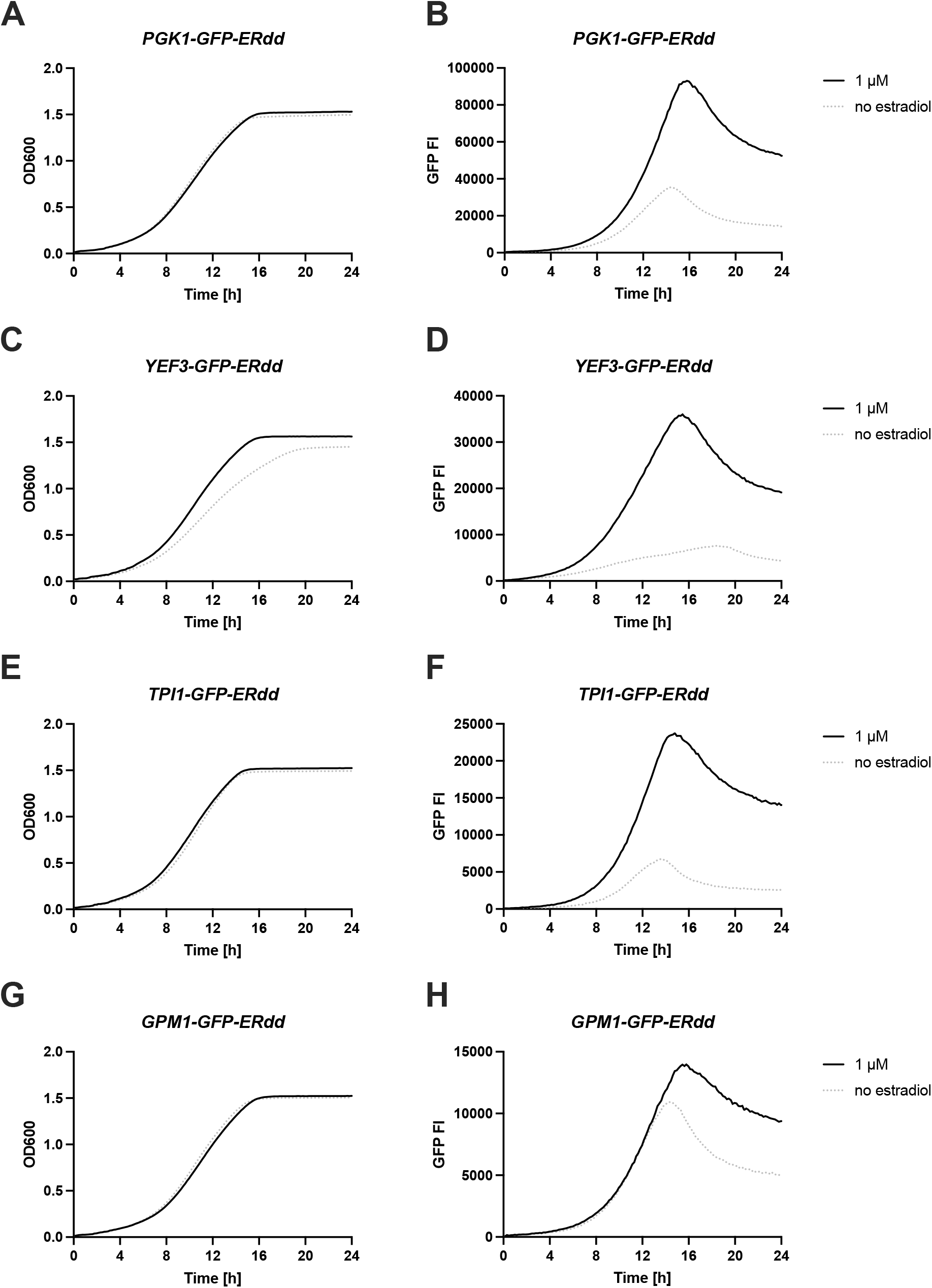
Liquid growth assays with recording of GFP fluorescence intensity of GFP-ERdd strains without and with 1 μM estradiol. Growth was monitored by measuring optical density at 600 nm (left column) and fluorescence intensity (right column) as measure for relative abundance of the essential GFP-ERdd fusion protein.

**Figure S2:**
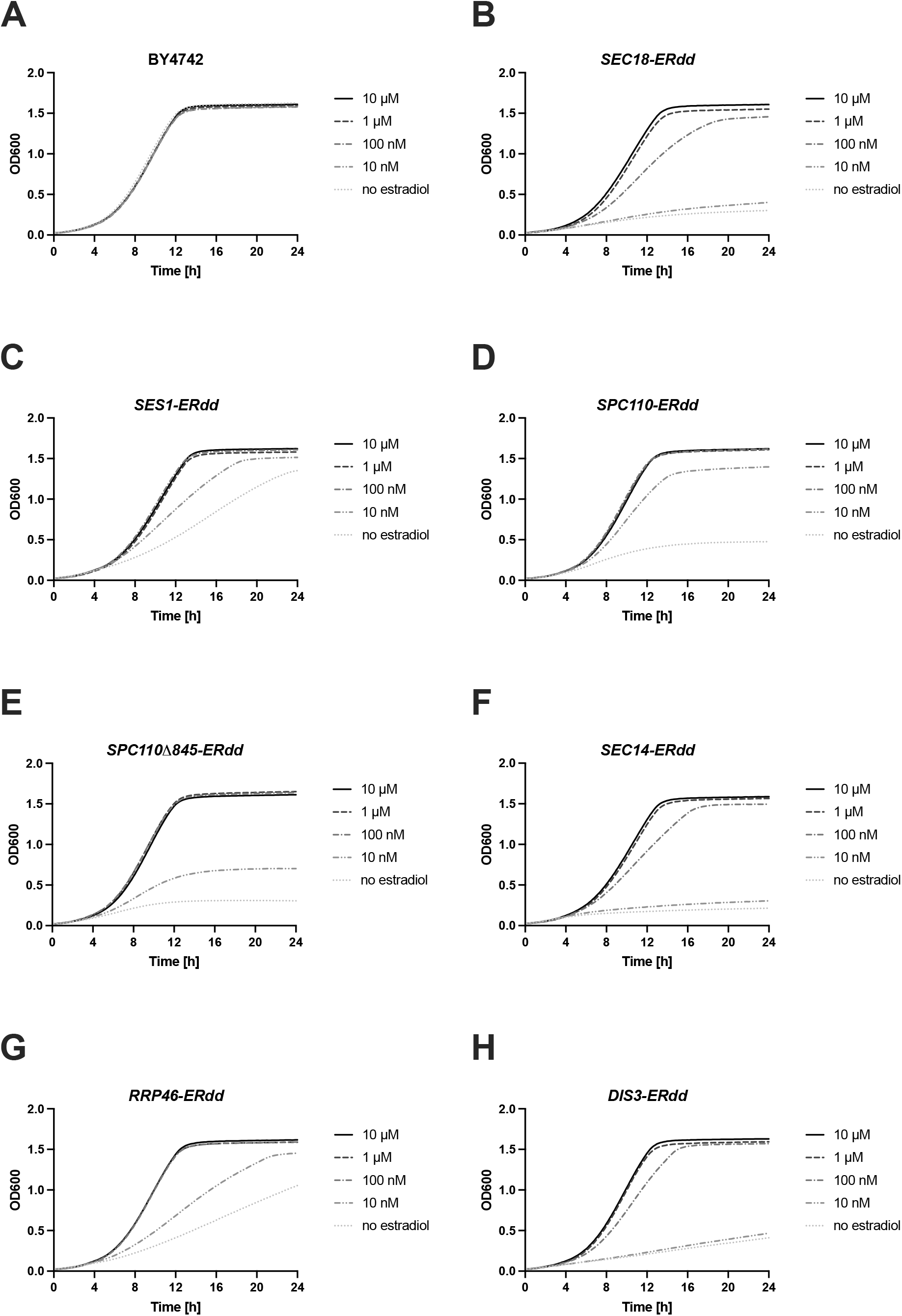
Liquid growth assays to assay estradiol dependence in wildtype BY4742 and singly ERdd-tagged strains.

**Figure S3:**
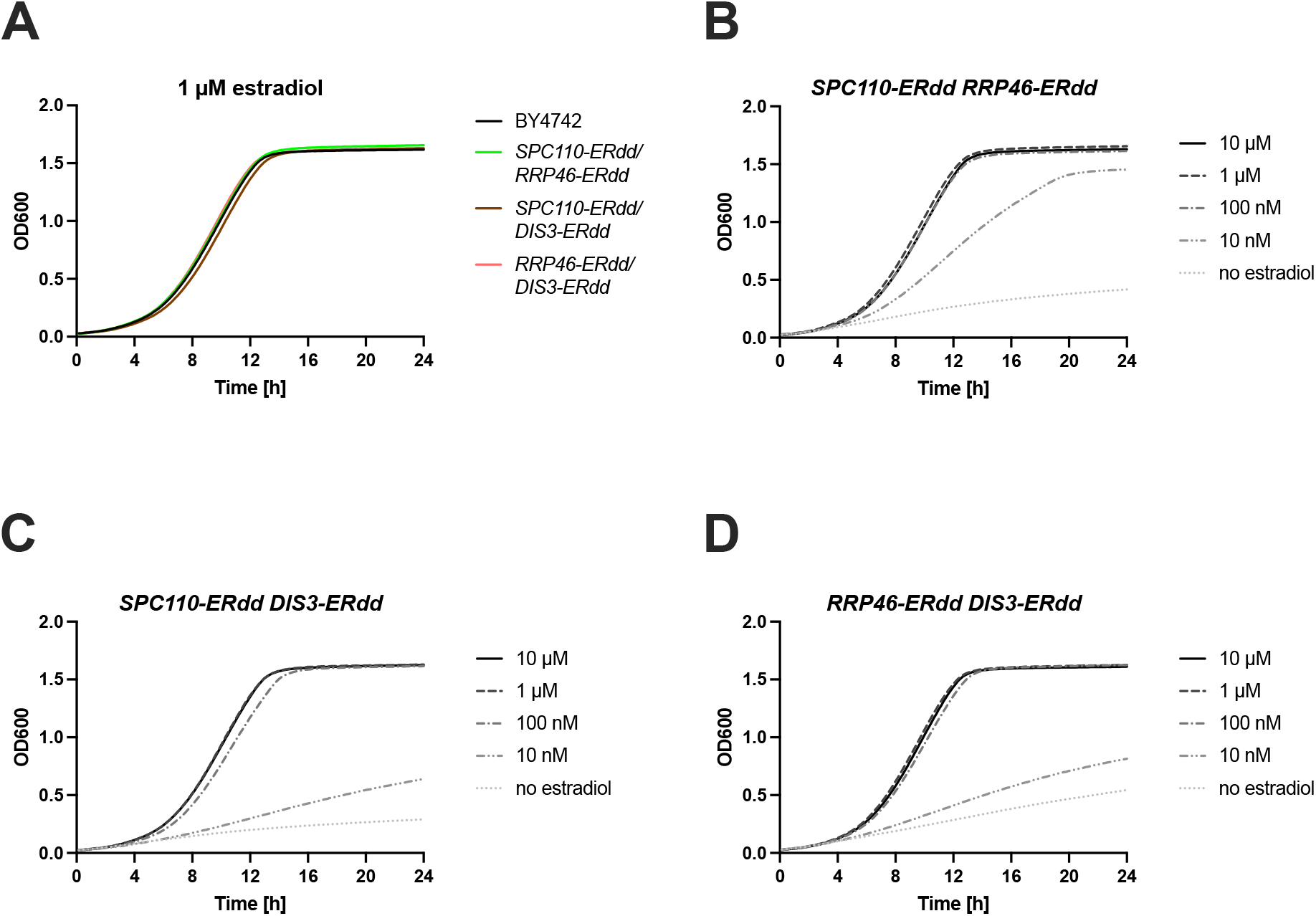
Liquid growth assays to assay estradiol dependence in strains with two select essential genes ERdd-tagged.

**Table S1:**
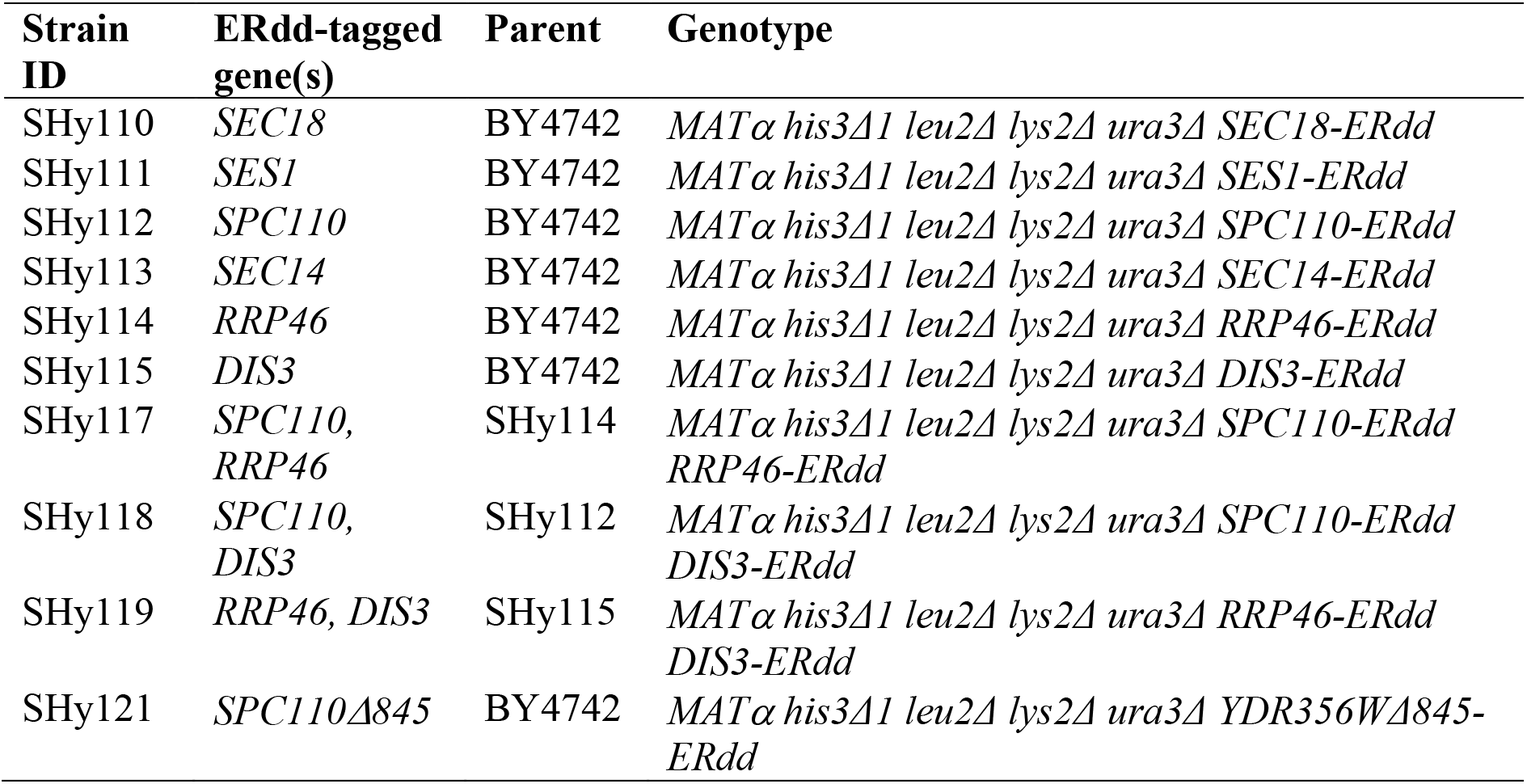
Generated direct ERdd fusion strains.

| Strain ID | ERdd-tagged gene(s) | Parent | Genotype |
| --- | --- | --- | --- |
| SHy110 | *SEC18* | BY4742 | *MATα his3Δ1 leu2Δ lys2Δ ura3Δ SEC18-ERdd* |
| SHy111 | *SES1* | BY4742 | *MATα his3Δ1 leu2Δ lys2Δ ura3Δ SES1-ERdd* |
| SHy112 | *SPC110* | BY4742 | *MATα his3Δ1 leu2Δ lys2Δ ura3Δ SPC110-ERdd* |
| SHy113 | *SEC14* | BY4742 | *MATα his3Δ1 leu2Δ lys2Δ ura3Δ SEC14-ERdd* |
| SHy114 | *RRP46* | BY4742 | *MATα his3Δ1 leu2Δ lys2Δ ura3Δ RRP46-ERdd* |
| SHy115 | *DIS3* | BY4742 | *MATα his3Δ1 leu2Δ lys2Δ ura3Δ DIS3-ERdd* |
| SHy117 | *SPC110, RRP46* | SHy114 | *MATα his3Δ1 leu2Δ lys2Δ ura3Δ SPC110-ERdd RRP46-ERdd* |
| SHy118 | *SPC110, DIS3* | SHy112 | *MATα his3Δ1 leu2Δ lys2Δ ura3Δ SPC110-ERdd DIS3-ERdd* |
| SHy119 | *RRP46, DIS3* | SHy115 | *MATα his3Δ1 leu2Δ lys2Δ ura3Δ RRP46-ERdd DIS3-ERdd* |
| SHy121 | *SPC110Δ845* | BY4742 | *MATα his3Δ1 leu2Δ lys2Δ ura3Δ YDR356WΔ845-ERdd* |

**Table S2:**
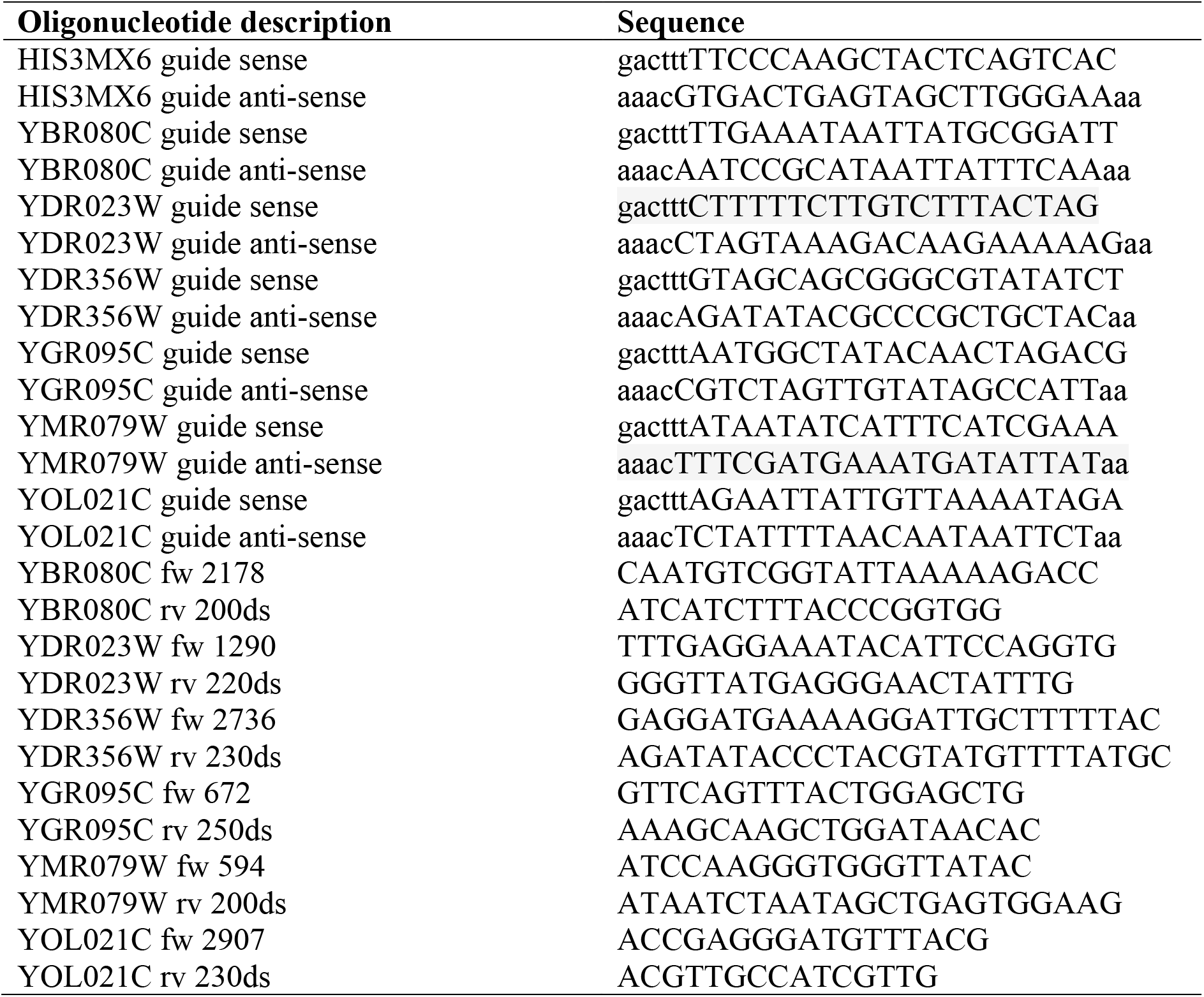
Oligonucleotides used in this study.

## Footnotes

1 https://rdrr.io/bioc/SLGI/man/essglist.html

